# Synthetic Genome Shuffling of Poxviruses through Yeast for Next-Generation Oncolytic Platforms

**DOI:** 10.64898/2026.03.06.710085

**Authors:** A. Agaoua, C. Rey, J. Hortelano, A-I Moro, B. Grellier, P. Erbs

## Abstract

Oncolytic viruses (OVs) are promising cancer therapeutics that selectively infect and lyse tumor cells while sparing normal tissues and stimulating antitumor immunity. However, their efficacy remains limited by suboptimal cytolytic activity and insufficient immune stimulation, highlighting the need for improved designs. Here, we introduce a synthetic virology platform leveraging transformation-associated recombination (TAR) in yeast to generate infectious chimeric poxviruses with enhanced therapeutic potential. Using TAR, we first cloned the Vaccinia virus (VACV) genome into a yeast plasmid and rescued it in human cancer cells. This plasmid was then co-transformed with Cowpox virus (CPXV) and Rabbitpox virus (RPXV) genomic DNA to promote recombination and create chimeric constructs. Subsequent rescue with Modified Vaccinia virus Ankara (MVA) yielded five infectious chimeric viruses. Phenotypic characterization revealed diverse plaque morphologies, comet-like spreading, and variable oncolytic activity across multiple cancer cell lines, indicating functional diversity arising from genome shuffling. Whole-genome sequencing confirmed recombination between VACV, CPXV, RPXV, and MVA. This study represents the first demonstration of TAR cloning for chimeric virus generation, establishing a versatile platform for designing next-generation oncolytic viruses.

## INTRODUCTION

Oncolytic viruses (OVs) are natural or genetically modified organisms capable of selectively infecting and lysing tumor cells while sparing normal tissues. In addition to their direct cytolytic effects, OVs can stimulate antitumor immune responses, further contributing to their therapeutic potential (Santos Apolonio *et al*, 2021). Compared to conventional cancer therapies, OVs offer the promise of reduced toxicity and enhanced efficacy, positioning them as attractive agents in cancer treatment (Vähä-Koskela *et al*, 2007; Russell *et al*, 2012). A diverse array of viruses has been explored for oncolytic applications, including adenoviruses, poxviruses, herpes simplex viruses, coxsackieviruses, polioviruses, measles virus, and Newcastle disease virus, several of them have reached the clinical trial stage. (Heise & Kirn, 2000; Dobrikova *et al*, 2008; Dingli *et al*, 2014; Fábián *et al*, 2007). To date, three OVs have received regulatory approval for cancer therapy (Macedo *et al*, 2020). Despite these advances, the clinical efficacy of current OVs remains limited by insufficient lytic activity and suboptimal immune stimulation, often resulting in incomplete tumor eradication (Macedo *et al*, 2020). This underscores the need for innovative strategies to develop more potent OVs. Directed evolution, which involves the selective adaptation of viruses within cancer cells through iterative infection cycles under stringent conditions, has been employed to enhance viral infectivity, spread, safety, and targeting (Kuhn *et al*, 2008; Shmulevitz *et al*, 2012). However, improvements achieved through this approach have generally been modest and limited to superficial phenotypic changes (Zainutdinov *et al*, 2019). An alternative and promising strategy is the generation of chimeric viruses (Ricordel *et al*, 2018a; El-Andaloussi *et al*, 2012). The Poxviridae family, comprising 18 genera and 46 known species capable of infecting vertebrates, includes several viruses with inherent oncolytic properties (Ricordel *et al*, 2018b). This diversity makes poxviruses excellent candidates for genome shuffling to create novel chimeric viruses. Leveraging the genetic diversity within this family, new viral species with enhanced oncolytic properties can be engineered. Notably, three chimeric poxviruses—deVV5, CF17, and CF33—have been developed for cancer therapy, demonstrating improved oncolytic activity compared to their parental strains (Ricordel *et al*, 2018a; O’Leary *et al*, 2018; Hammad *et al*, 2020). These chimeric viruses were generated via homologous recombination during co-infection of human cancer cell lines with multiple wild-type poxviruses. However, the combination of homologous recombination and selection in cell lines typically yields a limited number of chimeric viruses, primarily selected for replication efficiency. Other important oncolytic properties, such as extracellular enveloped virus (EEV) production or syncytia formation, may not be adequately represented (Law *et al*, 2006; Burton *et al*, 2019). Transformation-associated recombination (TAR) is a powerful cloning technique that exploits the homologous recombination machinery of yeast to assemble large DNA sequences, enabling the integration of up to 500 kb from multiple fragments in a single step (Kouprina & Larionov, 2008; Gibson *et al*, 2008). TAR has been successfully employed to clone complete viral genomes, including several large DNA viruses such as adenoviruses (32 kb), herpes simplex virus (152 kb), and human cytomegalovirus (235 kb) (Oldfield *et al*, 2017; Ketner *et al*, 1994; Vashee *et al*, 2017; Erbs *et al*, 2025). These cloned genomes can subsequently be used to reconstitute infectious viruses in appropriate host cells.

In our study, we established a poxvirus shuffling platform based on TAR cloning, enabling the generation of chimeric genomes independently of selection criteria imposed by cell culture. We first cloned the Vaccinia virus (VACV) genome into a yeast plasmid using TAR and regenerated infectious virus in human cancer cells. This plasmid was then used as a backbone for genome shuffling with other poxvirus genomes in *Saccharomyces cerevisiae*. The resulting chimeric viruses, recovered after rescue, exhibited a broad diversity of phenotypes and genotypes. To our knowledge, this is the first demonstration of TAR cloning for the generation of chimeric poxvirus genomes, providing a versatile platform for the rational design of next-generation oncolytic viruses.

## RESULTS

### TAR cloning strategy and virus rescue

The schematic overview of the construction of an infectious clone of VACV is presented in Figure 1. The VACV genome spans approximately 191 kb (Scott J Goebel *et al*, 1990), comprising a unique central genomic region of approximately 165 kb that encodes around 247 putative open reading frames (ORFs). Flanking this region are the 5′ and 3′ inverted terminal repeats (ITRs), each comprising three distinct structural elements: i) coding ITRs (ITRc), approximately 8 kb in length, containing ∼13 conserved ORFs; ii) repeat regions (RRs), approximately 5 kb of short tandem repeats; iii) hairpin structures (Hp), approximately 0.15 kb of folded single-stranded DNA located at each extremity of the genome. In *S. cerevisiae*, the presence of both 5′ and 3′ inverted terminal repeats (ITRs) within a single plasmid induces significant instability, most likely due to homologous recombination between the ITR sequences. This recombination event leads to the deletion of the intervening genomic region, resulting in a truncated plasmid that lacks the core of the viral genome. To circumvent this issue, the genome of the VACV *Copenhagen* strain was assembled into a yeast plasmid containing the core genome, the 5′ ITRc and the 5’RR region, thereby improving plasmid stability.

**Figure 1:**
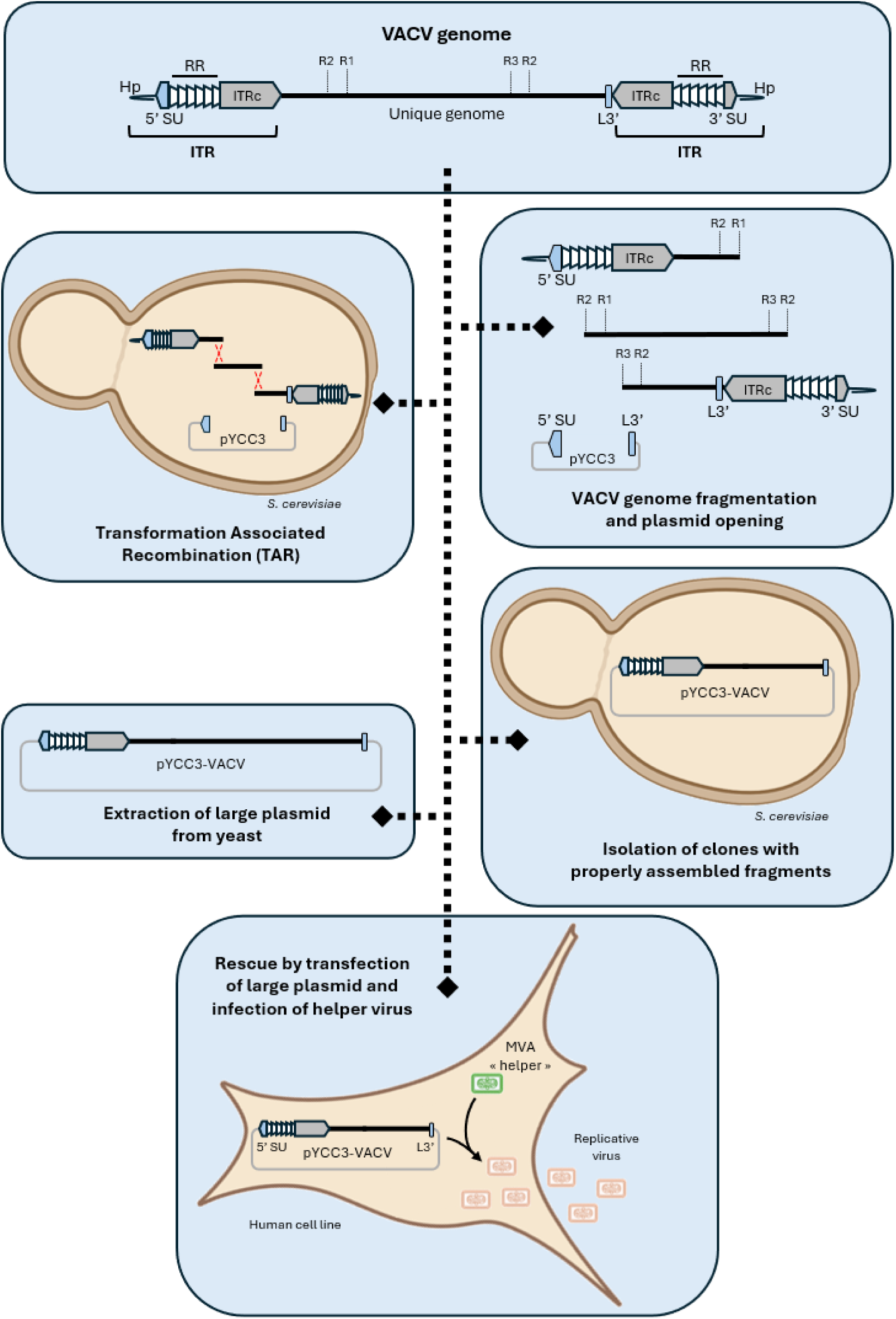
Strategy of the VACV genome assembly and virus rescue. The VACV viral genome is divided into three overlapping fragments, which are assembled into a single plasmid through homologous recombination of shared sequences in yeast. The assembled plasmids are then purified and transfected into cells in the presence of a non-replicative helper virus, enabling the production of infectious viral particles. Abbreviations: ITR, *Inverted Terminal Repeat*; ITRc, *Coding Inverted Terminal Repeat*; RR, *Repeat Region*; Hp, *Hairpin*; R1, R2, R3, positions of restriction sites used to generate the three overlapping fragments; SU, *Small Unique region*; L3’, terminal region of the last gene at the 3′ end of the unique genome.

Yeast centromeric and auxotrophic selection markers were introduced into an *E. coli* single-copy vector using conventional molecular biology techniques (Wittek & Moss, 1980). The vector, pYCC3, was engineered to include hook sequences corresponding to the 5′ SU and L3′ regions of the VACV genome. The 5′ SU hook excludes the 5′ Hp region, which consists of single-stranded folded DNA that cannot be accommodated in a circular double-stranded plasmid. The L3′ hook is positioned at the boundary of the 3′ end of the unique genomic region, within the B19R gene, thereby excluding the 3′ ITRc, RR, and Hp regions from recombination. TAR technology exploits the high-efficiency homologous recombination machinery of yeast to assemble multiple overlapping DNA fragments. In this strategy, three overlapping VACV fragments, designated R1, R2, and R3, were co-transformed with the pYCC3 vector into *S. cerevisiae*. Following successful assembly, the recombinant plasmid was extracted and transfected into human cells pre-infected with a helper virus. Because poxvirus DNA alone is not infectious (Moss, 2013), the Modified Vaccinia virus Ankara (MVA) strain was used as a helper virus. MVA is replication-deficient in human cells but provides essential viral proteins and complements missing genomic regions, enabling the rescue and replication of the recombinant VACV genome (P. Erbs, personal communication).

### VACV genome assembly via TAR Cloning

The VACV genome was fragmented using three restriction enzyme digestions, generating ∼70 kb fragments with overlapping sequences to facilitate homologous recombination. These three VACV fragments, along with the linearized pYCC3 vector, were co-transformed into *S. cerevisiae* spheroplasts using the TAR method. Of the 188 TRP⁺ yeast clones screened, 18 were tested positive for the presence of viral DNA in the first round of PCR targeting VACV genomic sequences. A second round of PCR, targeting each of the three VACV genome fragments individually, was performed on seven of these clones. All seven were confirmed to contain the complete set of fragments (Figure 2A). DNA extracts from two representative yeast clones, C1 and C2, which carried the expected ∼188 kb construct, were electroporated into *E. coli*. Although transformation efficiency is typically low for large constructs, several bacterial clones were successfully obtained. Plasmid DNA from these clones, designated pYCC3-VACV_C1 (noted C1) and pYCC3-VACV_C2 (noted C2), was validated by restriction enzyme profiling (Figure 2B). Subsequent analysis using Oxford Nanopore Technology sequencing confirmed the integrity of the assembled VACV genome. The viral genome was precisely inserted between the 5′ SU and L3′ hook sequences within the pYCC3 vector, with no mutations detected across the assembled region. Notably, the 5′ RR region was found to be shorter in both clones compared to the reference genome. However, this region is known to be hypervariable, and its truncation does not interfere with downstream applications or viral functionality (Wittek & Moss, 1980; Moss *et al*, 1981).

**Figure 2:**
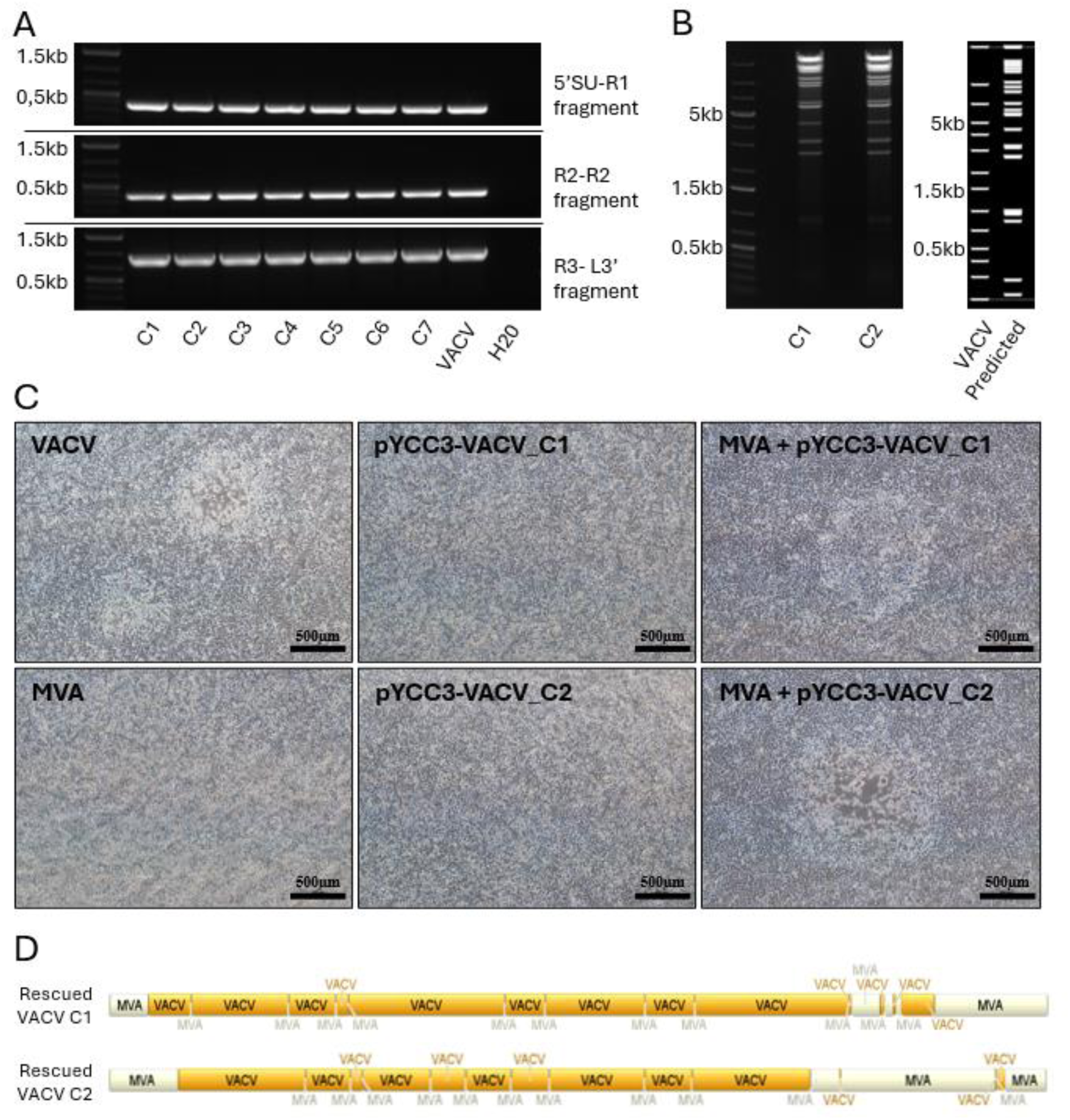
Analysis of isolated clones from TAR to rescue. (A) PCR analysis of the three VACV genomic fragments transformed into yeast, performed on pre-selected TAR-derived clones. (B) Restriction enzyme analysis of plasmid DNA from clones C1 and C2 following transformation into *E. coli* and plasmid purification. (C) Brightfield microscopy images of plaque assays on A549 cells infected with MVA, VACV, the rescue material from cells transfected with pYCC3-VACV_C1 (clone C1), the rescue material from cells transfected with pYCC3-VACV_C2 (clone C2), as well as rescue samples from MVA-infected cells transfected with pYCC3-VACV_C1 (MVA + pYCC3-VACV_C1) or pYCC3-VACV_C2 (MVA + pYCC3-VACV_C2). A549 cells were seeded at 5 × 10⁵ cells per well in 6-well plates and infected at a multiplicity of infection (MOI) of 10⁻⁵. Images were acquired 48 h post-infection at 4× magnification using brightfield microscopy. (D) Distribution of parental viral genomes within isolated clones obtained from rescue experiments in MVA-infected cells transfected with pYCC3-VACV_C1 or pYCC3-VACV_C2, based on deep sequencing analysis.

### Rescue and Characterization of Synthetic Infectious VACVs

To verify that the VACV genome—lacking the 3′ ITRc, 3′ RR, and Hp regions—inserted into the pYCC3 vector could be complemented to generate infectious viral particles, a rescue experiment was performed (Figure 2C). Human A549 cells were first infected with MVA, followed by transfection with either pYCC3-VACV_C1 or pYCC3-VACV_C2, both plasmids previously isolated from *S. cerevisiae*. No cytopathic effect was observed in cells infected with MVA alone or in cells transfected with the plasmids alone. In contrast, cell lysis was clearly observed in MVA-infected cells transfected with either pYCC3-VACV_C1 or pYCC3-VACV_C2, forming lysis plaques similar to those induced by wild-type VACV. Two plaques from the rescued viruses were isolated and subjected to genome sequencing (Figure 2D). Sequencing confirmed the presence of VACV genomes derived from the pYCC3-VACV constructs. As expected, the missing genomic regions were complemented by sequences from the MVA genome. Recombination between VACV and MVA occurred differently in the two clones, although both exhibited similar overall genomic profiles. In the 5′ region, recombination restored the Hp structure using MVA-derived sequences. In the 3′ region, the 3′ ITRc, RR, and Hp were also complemented by MVA. Additionally, small MVA-derived insertions were detected throughout the rescued VACV genomes. These results demonstrate that the pYCC3-VACV plasmid can be used for regenerating infectious VACV particles when complemented by MVA in trans.

### TAR shuffling strategy

The schematic overview for constructing infectious chimeric poxviruses using TAR is presented in Figure 3. To generate shuffled chimeric genomes, the pYCC3-VACV plasmid, along with genomic DNA from Cowpox virus (CPXV) and Rabbitpox virus (RPXV), was co-transformed into *S. cerevisiae*. To enhance transformation efficiency and promote homologous recombination, the viral genomes were digested with restriction enzymes to produce fragments ranging from 10 to 40 kb (Kouprina *et al*, 2021). The restriction enzyme used for pYCC3-VACV was specifically selected to avoid cleavage within the vector backbone, thereby preserving its structural integrity during recombination. Two TRP⁺ clones were obtained when *S. cerevisiae* was transformed with digested pYCC3-VACV alone and none with digested CPXV and RPXV genomic DNA alone. In contrast, 42 TRP⁺ clones were recovered when *S. cerevisiae* was co-transformed with digested pYCC3-VACV and digested genomic DNA from CPXV and RPXV, indicating successful recombination events. To evaluate the formation of chimeric viral DNA constructs, plasmids were extracted from ten TRP⁺ yeast clones and subjected to a rescue assay using MVA as a helper virus in HeLa cells. As previously described, no cytopathic effect was observed in cells infected with MVA alone or transfected with plasmids alone. However, in MVA-infected cells transfected with plasmids isolated from *S. cerevisiae*, five clones successfully produced infectious viral particles, confirming the feasibility of TAR-based shuffling for synthetic poxvirus engineering.

**Figure 3:**
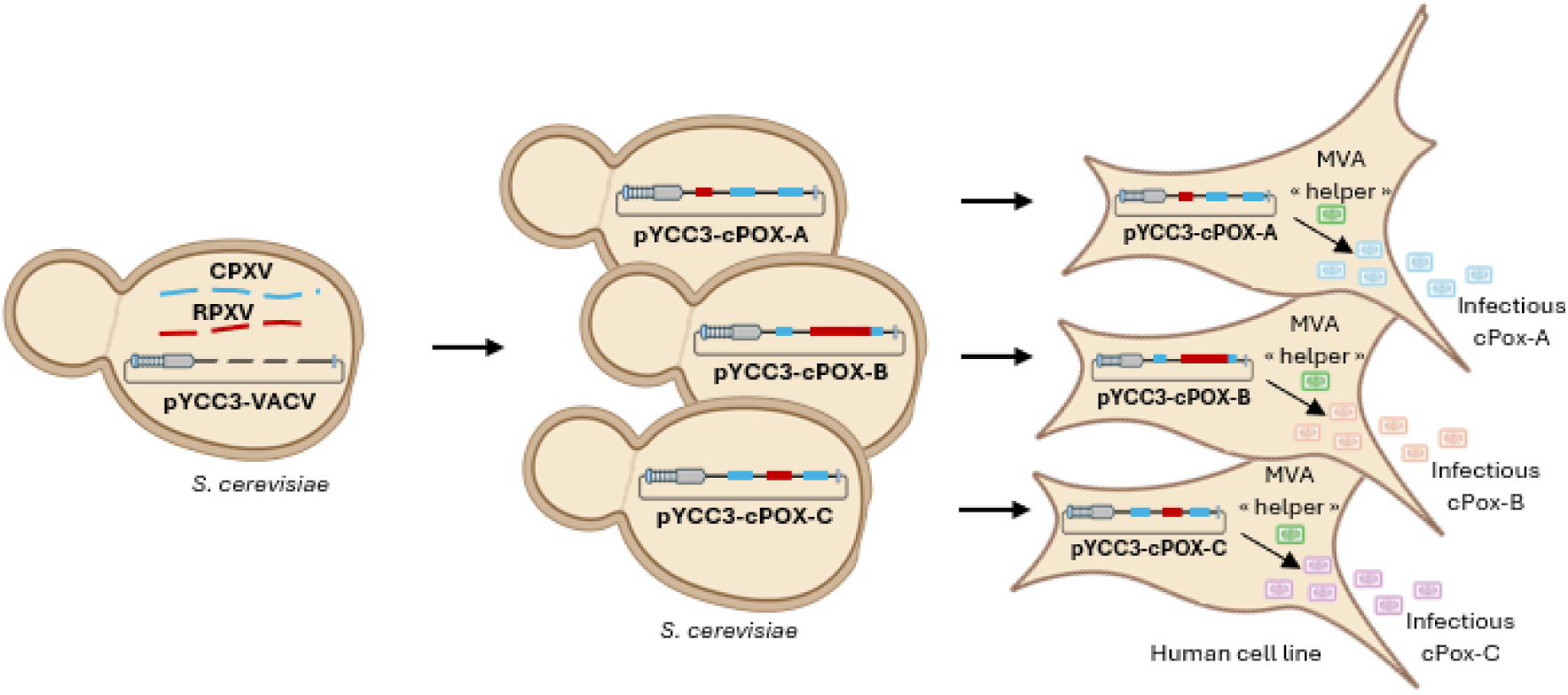
TAR-based shuffling of CPXV and RPXV genomes using the pYCC3-VACV plasmid. Yeast cells were co-transformed with fragmented pYCC3-VACV plasmid DNA together with CPXV and RPXV genomic DNA. Each resulting yeast colony is expected to contain a distinct chimeric poxvirus sequence (pYCC3-cPOX) generated through homologous recombination. DNA extracted from individual colonies was subsequently transfected into human cells pre-infected with the helper virus MVA, enabling the production of chimeric infectious viral particles.

### Characterization of synthetic infectious chimeric poxviruses

Plaque morphology and syncytia formation were evaluated in HeLa cells (Figure 4A), and viral dissemination was assessed by comet assay in A549 cells (Figure 4B) for the parental viruses (excluding MVA, which does not replicate or form plaques in human cells) and the cPOX variants. Among the parental strains, VACV and CPXV produced regular plaques without syncytia or comet formation, whereas RPXV generated syncytial plaques in HeLa cells and exhibited a spreading phenotype in A549 cells. The cPOX01-02 clone displayed VACV-like lytic plaques in both HeLa and A549 cells but showed limited dissemination. cPOX02-03 formed syncytial plaques with irregular contours in HeLa cells and demonstrated low spreading in A549 cells. cPOX04-12 produced small RPXV-like plaques in HeLa cells and extensive comet formation in A549 cells. cPOX05-14 exhibited irregular lysis with small syncytia in HeLa cells and lacked comet formation in A549 cells. Finally, cPOX06-18 generated small, dispersed plaques with syncytia in both HeLa and A549 cells.

**Figure 4:**
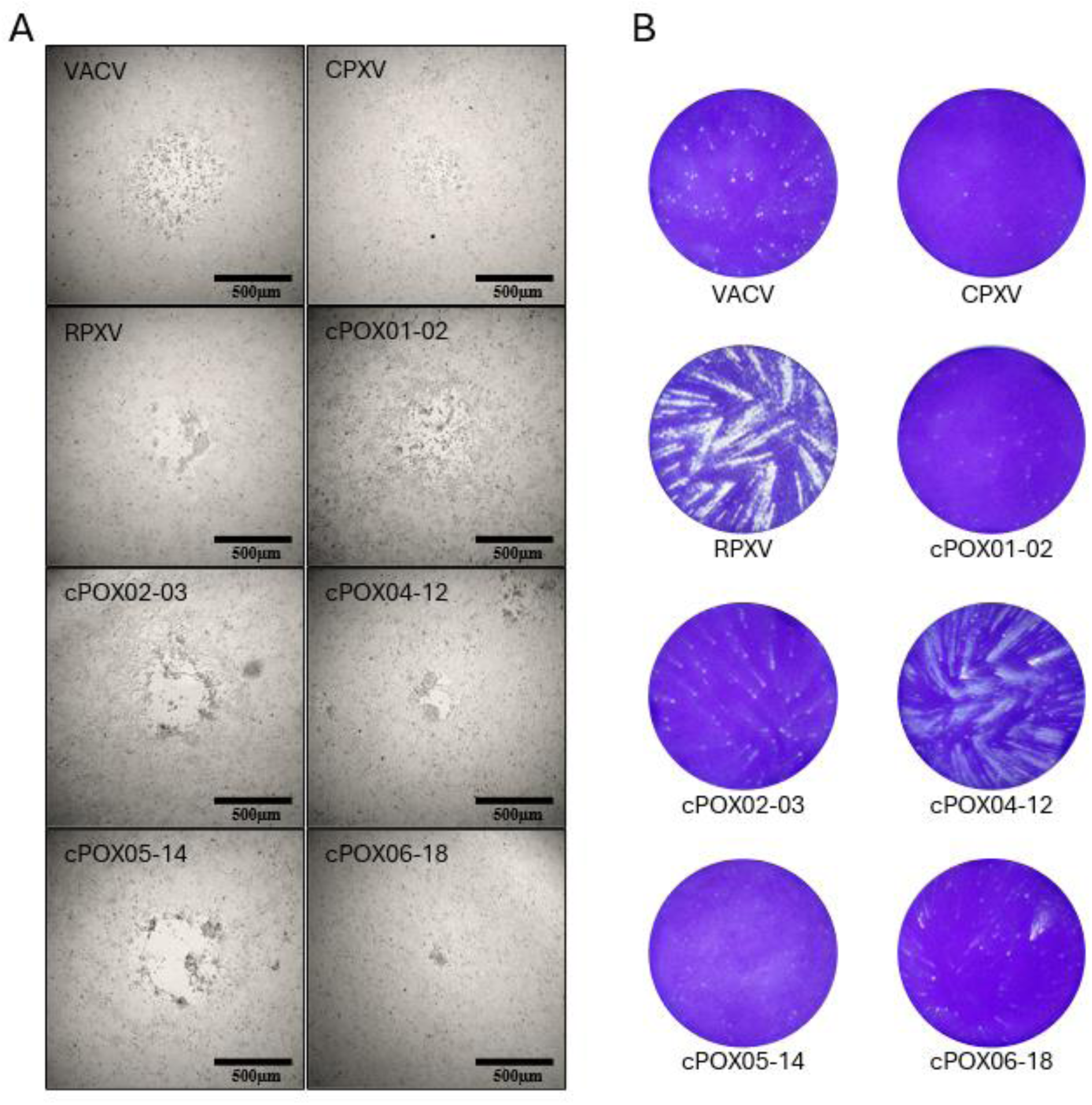
Plaque morphology of chimeric viruses. (A) Brightfield microscopy images of plaque assays performed on HeLa cells infected with parental viruses (VACV, CPXV, RPXV) and with chimeric viruses rescued from TAR-shuffling clones (cPOX01-02, cPOX02-03, cPOX04-12, cPOX05-14, and cPOX06-18). HeLa cells were seeded at 5 × 10⁵ cells per well in 6-well plates and infected at a multiplicity of infection (MOI) of 10⁻⁵. Images were acquired 48 h post-infection at 4× magnification. (B) Representative comet-shaped plaques observed in A549 cells. A549 monolayers were infected with the indicated viruses, incubated for 48 h, and stained with crystal violet to visualize comet formation, reflecting enhanced viral spread

Oncolytic potency was evaluated across four human tumor cell lines (A549, HeLa, HCT116, and MIA PaCa-2) by determining half-maximal effective concentrations (EC_50_) values following infection with serial dilutions of each virus (Figure 5). Among the parental strains, VACV exhibited strong activity with EC_50_ values ranging from 1.2 × 10^-2^ to 2.7 × 10^-2^ MOI, whereas CPXV and the non propagative MVA were markedly less potent (EC_50_ > 0.3 MOI). RPXV showed intermediate activity, particularly in HeLa cells (4.7 × 10^-2^ MOI). Notably, several cPOX variants demonstrated enhanced potency compared to VACV, with cPOX01-02 and cPOX04-12 achieving the lowest EC_50_ values in A549 (9 × 10^-3^ MOI) and HCT116 (1.9 × 10^-3^ MOI), respectively. These findings indicate that specific modifications in cPOX variants significantly improve oncolytic activity, particularly in colorectal and lung cancer cell lines.

**Figure 5.**
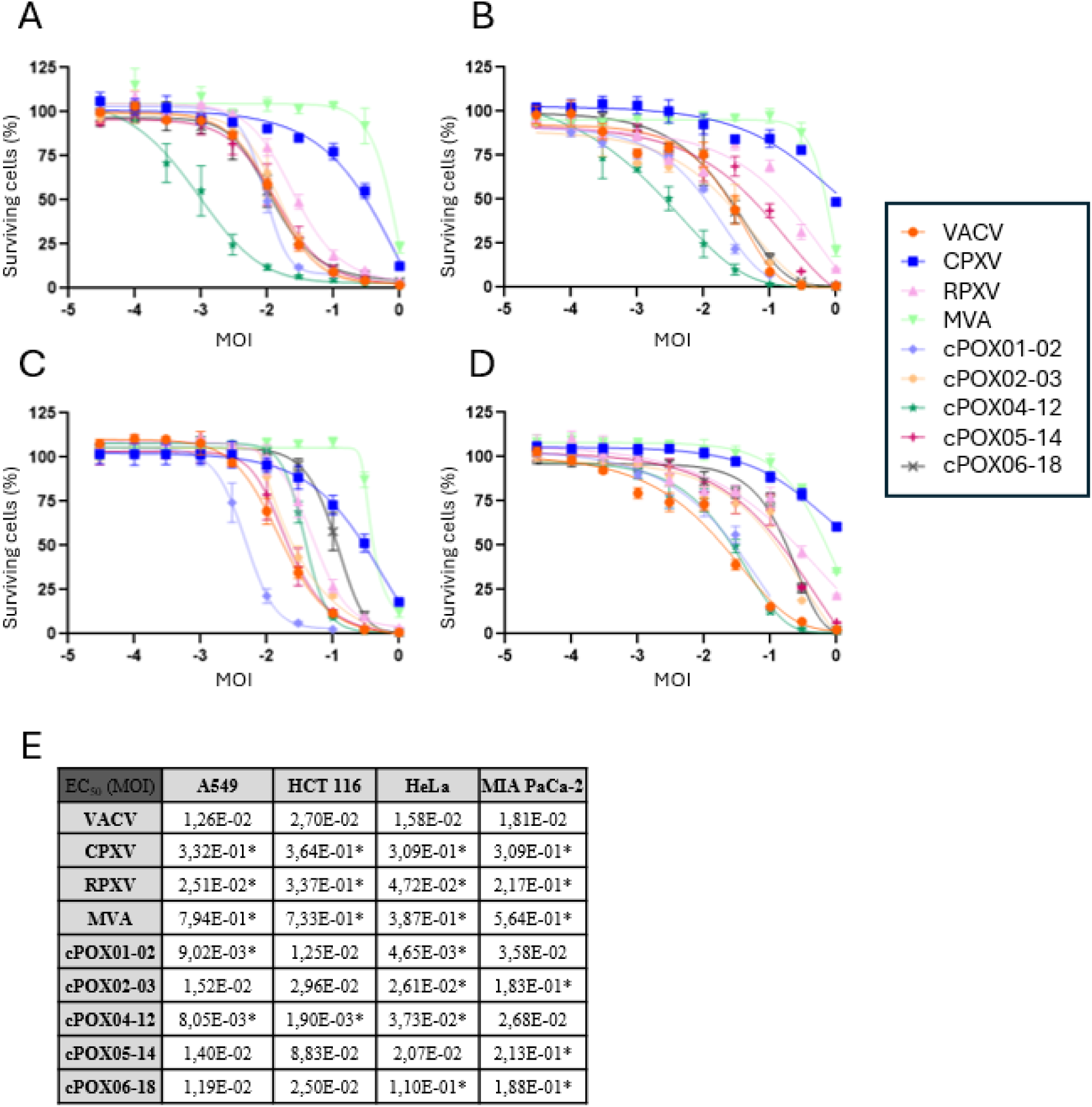
Oncolytic activity across a panel of human tumor cell lines. (A–D) A549 (A), HCT 116 (B), HeLa (C), and MIA PaCa-2 (D) cells were infected with serial dilutions of each virus. Cell viability was assessed 4 days post-infection using the CellTiter-Blue assay and normalized to uninfected controls. Data represent the mean ± SD from three independent experiments. (E) EC₅₀ values for each virus in each cell line were calculated by fitting a sigmoidal dose–response curve to the viability data shown in panels A–D. Statistical significance was determined relative to the VACV group using one-way ANOVA: *p* < 0.05 (*).

All tested viruses replicated in A549 cells (Figure 6A), with VACV showing the highest amplification (∼2 x 10^5^ fold). CPXV and RPXV exhibited intermediate amplification (∼10^5^), while cPOX04-12 reached levels comparable to VACV. Other chimeric strains displayed variable replication, with cPOX01-02 being the least efficient (∼10^4^ fold). Analysis of EEV production revealed marked differences among strains (Figure 6B-C). VACV and CPXV produced minimal EEV (< 0.2 %), RPXV reached ∼4 %, and cPOX04-12 showed the highest proportion (∼9 %). These EEV percentages are consistent with comet formation observed, with cPOX04-12 exhibiting extensive spreading. Together, these data indicate that cPOX04-12 combines strong replication capacity with enhanced EEV release, features that may favor efficient cell-to-cell spread.

**Figure 6:**
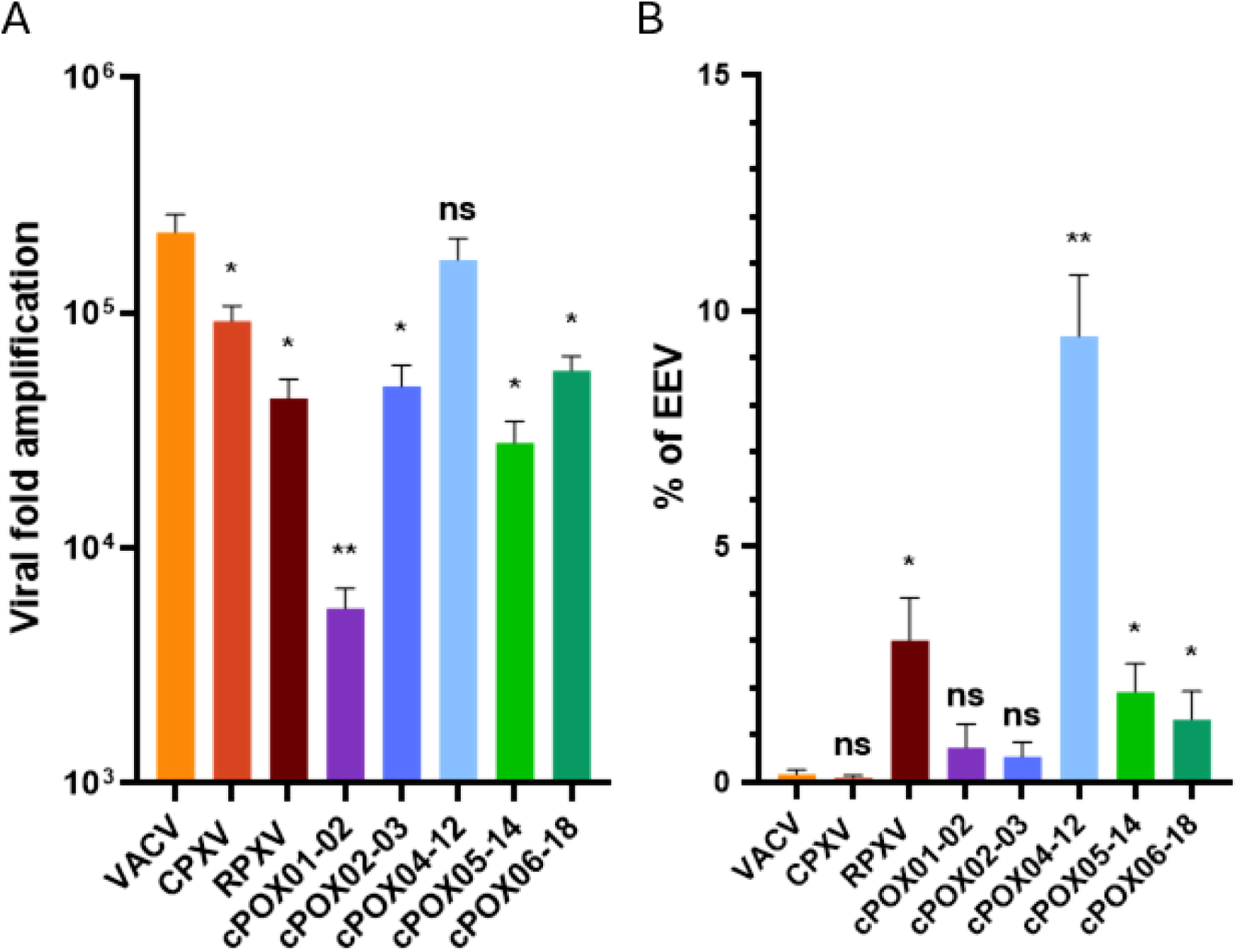
Replication and early production of EEV in tumor cells. (A) Viral amplification in A549 cells. Monolayers were infected at a MOI of 10⁻³. At 3 days post-infection, both supernatants and cells were collected, and viral progeny were quantified by plaque assay. Results are expressed as fold amplification, calculated as the output/input ratio. Data represent mean ± SD from three independent experiments. Statistical significance was assessed against the VACV group using one-way ANOVA: not significant (ns), *p* < 0.05 (*), *p* < 0.01 (**). (B) Ratio of EEV to total progeny virus (IMV + EEV). A549 cell monolayers were infected with the indicated viruses at a MOI of 0.1. Viral titers were determined for EEV (supernatant only) and total progeny virus (IMV + EEV; combined supernatants and cells). Data represent mean ± SD from three independent experiments. Statistical significance was assessed against the VACV group using one-way ANOVA: not significant (ns), *p* < 0.05 (*), *p* < 0.01 (**).

### Sequence analysis

To identify recombination events that occurred during the TAR cloning and rescue process, the five clones were subjected to whole-genome sequencing using nanopore sequencing technology (Figure 7). For each clone, unique recombination events involving sequences from VACV, CPXV, RPXV, and MVA were identified. Sequenced genome sizes for the different clones ranged from 150 kb to 187 kb. However, the inverted terminal repeats (ITR) were only partially covered by sequencing, preventing any conclusions regarding their structure, function, or potential variability. The presence of CPXV and RPXV genomic segments confirms that homologous recombination occurred in *S. cerevisiae* during co-transformation with the pYCC3-VACV vector. Additionally, sequences derived from the MVA genome were found in the rescued viruses, indicating that recombination between the chimeric plasmid and MVA took place during the rescue step in human cells. These findings demonstrate that infectious chimeric viruses can be generated from four distinct parental genomes, highlighting the versatility of the TAR shuffling strategy for synthetic virology and oncolytic virus development. Moreover, the observed phenotypic diversity and variable oncolytic potential among the clones underscore the utility of this approach for engineering novel viral therapeutics.

**Figure 7:**
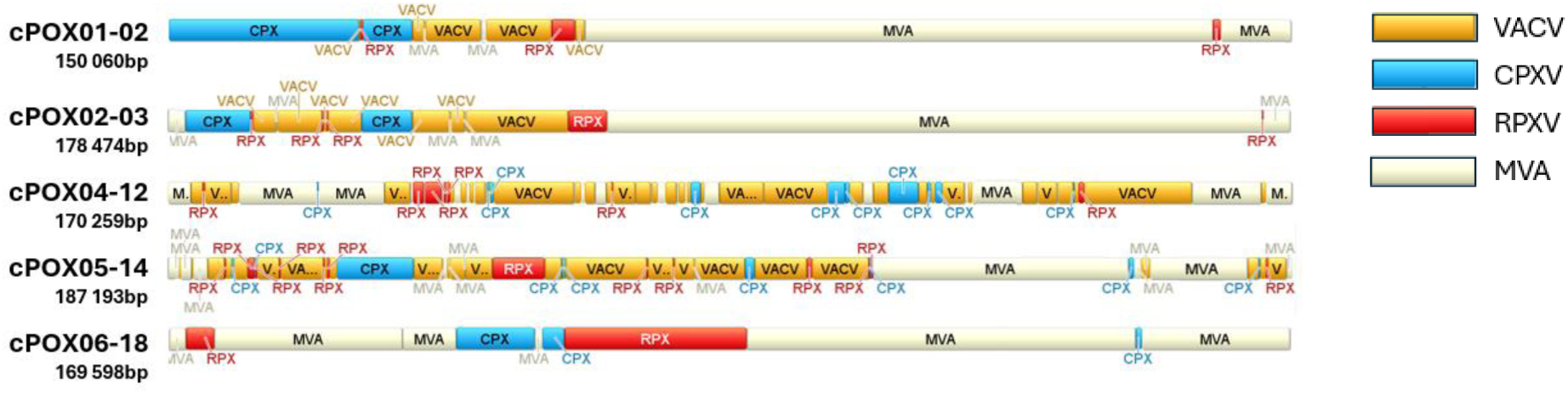
Genomic map of cPOX viruses showing the origin of genomic segments. Genomic regions are color-coded according to their respective parental viruses. Each annotation highlights the longest contiguous segment displaying the highest percentage of sequence identity with the corresponding parental genome. Genome sizes are indicated beneath each virus name.

## DISCUSSION

Chimeric virus generation represents a powerful strategy for engineering novel OVs with enhanced therapeutic properties (El-Andaloussi *et al*, 2012; Ricordel *et al*, 2018a; Kuhn *et al*, 2008; O’Leary *et al*, 2018). By combining genetic elements from different viral backbones, this approach enables the rational design of recombinant viruses that integrate desirable features such as improved tumor selectivity, enhanced immunogenicity, and optimized replication kinetics. Such chimeric constructs can overcome limitations inherent to individual parental strains and offer a flexible platform for tailoring OVs to specific clinical contexts. In this study, we applied TAR technology to generate chimeric oncolytic poxviruses.

TAR technology, originally developed for cloning large genomic fragments in yeast, has emerged as a powerful tool for reconstructing complete viral genomes directly from DNA or clinical samples (Larionov *et al*, 1996; Oldfield *et al*, 2017; Gibson *et al*, 2040; Labroussaa *et al*, 2021). This method exploits the high efficiency of homologous recombination in *S. cerevisiae*, allowing the assembly of large DNA molecules from overlapping fragments or even from fragmented genomic DNA. It has enabled the successful cloning of viral genomes across a wide size range, from the 10 kb Potyvirus to the 235 kb Human Cytomegalovirus (Desbiez *et al*, 2012; Brown *et al*, 2017). More recently, TAR has been used to reconstruct RNA viruses such as SARS-CoV-2 allowing rapid generation of infectious clones for functional studies and vaccine development (Thi Nhu Thao *et al*, 2020). Additionally, TAR has facilitated the assembly of large DNA viruses such as ascoviruses and baculoviruses, further demonstrating its broad applicability in synthetic virology (Zaghloul *et al*, 2025; Shang *et al*, 2017). Despite these advances, no complete poxvirus genome had been cloned into yeast prior to this study, likely due to the structural complexity and large size of poxvirus genomes, which include covalently closed hairpin termini and extensive ITRs, which pose challenges for stable maintenance in yeast artificial chromosomes. Our study addresses this limitation by applying TAR technology to assemble a near-complete VACV genome and to generate chimeric oncolytic poxviruses. We first used TAR to isolate and assemble most of the vaccinia virus (VACV) genome, deliberately excluding the unstable 5′ terminal hairpin and the complete 3′ inverted terminal repeat (ITR) region. This approach enabled the reconstruction of 175 kb of the 191 kb VACV genome. To complement the missing essential terminal sequences, previously described rescue approaches using Fowlpox virus (FPV) or Shope Fibroma virus (SFV) were considered. However, these viruses failed to support recombination with VACV in cell lines, likely due to the genetic divergence between avipoxvirus, leporipoxvirus, and orthopoxvirus (Scheiflinger *et al*, 1992; Yao & Evans, 2003; Qin & Evans, 2014). To overcome this limitation, we performed virus reactivation using MVA, an attenuated orthopoxvirus that, despite being replication-deficient in human cells, shares sufficient genomic homology with VACV to enable recombination and successful rescue. Following reactivation with MVA, the synthetic VACV particles exhibited the expected genomic architecture, with MVA-derived sequences incorporated at both the 5′ and 3′ termini. This outcome was consistent with homologous recombination mechanisms occurring within cytoplasmic viral factories, where the synthetic VACV genome interacts with the MVA helper virus. The incorporation of MVA terminal sequences underscores the importance of selecting a helper virus with adequate genomic compatibility to ensure efficient rescue and structural integrity of the recombinant virus. Importantly, the synthetic VACV constructs generated through this approach were fully replication-competent in human tumor cell lines and displayed phenotypic characteristics indistinguishable from wild-type VACV, including replication kinetics, cytopathic effects, and plaque morphology. These findings confirm that the TAR-based assembly strategy, despite omitting terminal regions, preserves the essential functional integrity of the virus when complemented appropriately. A second round of TAR cloning was performed to integrate additional poxvirus genomes into the previously constructed plasmid containing the near-complete VACV genome. Fragmented genomes from CPXV and RPXV were co-transformed with pYCC3-VACV into *S. cerevisiae*, enabling *in vivo* homologous recombination and the generation of novel chimeric genomes. Due to the technical challenge of sequencing dozens of 180 kb single-copy plasmids directly in *S. cerevisiae*, ten clones were selected for direct rescue. Remarkably, half of these yeast clones yielded infectious viruses upon reactivation. Sequencing of the rescued viruses revealed the presence of sequences derived from VACV, CPXV, and RPXV, confirming that homologous recombination occurred in *S. cerevisiae* and successfully generated chimeric viral genomes. These results demonstrate that TAR cloning can efficiently produce random chimeric constructs, offering a high degree of genomic diversity. Additionally, large DNA fragments derived from the MVA genome were detected in the sequenced chimeric constructs, suggesting that further recombination events occurred during the rescue process. This additional layer of recombination contributes to the overall genomic heterogeneity of the resulting viral population, further expanding the diversity generated through TAR cloning. Non-infectious clones may result from incomplete genome assembly during TAR cloning or from the generation of non-viable chimeric viruses that fail to replicate in the rescue cell line. The comparative analysis of chimeric clones revealed substantial heterogeneity in lysis morphology, viral replication and oncolytic activity across various human cancer cell lines. While clone, such as cPOX02-03 exhibited lysis pattern resembling none of the parental viruses with large syncytia formation. Others like cPOX05-14 and cPOX06-18 displayed atypical lysis phenotypes and reduced activity. More particularly, cPOX01-02 and cPOX04-12 demonstrated distinct functional profiles: cPOX01-02 outperformed VACV in HeLa cells despite displaying a VACV-like phenotype, while cPOX04-12 outperformed VACV in A549 and HCT 116 cells and additionally exhibited pronounced comet-like spreading. These findings underscore the functional diversity resulting from recombination-driven genome mosaicism and highlight the need for individual clone characterization to identify candidates with optimal therapeutic potential. Plaque and comet assays revealed notable phenotypic differences between the isolated chimeric clones and their parental viruses. In HeLa cells, infection resulted in distinct lysis patterns, with large syncytia in cPOX02-03 and smaller one observed in clones cPOX05-14 and cPOX06-18. Syncytia formation in poxvirus-infected cells has been primarily attributed to the A56R gene, which encodes a fusion-inhibitory protein. This protein, also known as the hemagglutinin, forms a complex with K2L, a serine protease inhibitor, on the surface of infected cells. Together, they prevent cell-cell fusion, thereby inhibiting syncytia formation (Turner & Moyer, 1995). When A56R is mutated or deleted, this inhibitory mechanism is lost, allowing the viral fusion machinery to mediate the fusion of infected cells with neighboring cells, leading to the formation of multinucleated syncytia (de Haven *et al*, 2011). However, all sequenced clones in this study carry intact A56R sequences derived from either MVA or VACV, suggesting that the syncytia observed in these clones are mediated by alternative genetic determinants yet to be identified. Comet formation, a distinct cytopathic effect observed in poxvirus-infected monolayers, is closely associated with the release of EEV (Smith & Law, 2004). EEV particles are enveloped in a double membrane derived from the host cell and are released prior to cell lysis, enabling long-range dissemination of the virus. This phenotype is particularly relevant in the context of viral spread and immune evasion(Vanderplasschen *et al*, 1998). Comet formation was minimal in all chimeric clones except cPOX04-12, which displayed prominent comet-like structures in A549 cells, associated with a markedly higher proportion of EEV, showing an EEV production rate approximately 50-fold greater than that of VACV. The A34R gene is the only poxvirus gene currently known to regulate EEV formation, with a single mutation at residue K151 (K151E) shown to enhance EEV production up to 40-fold in the Vaccinia strain (Mcintosh & Smith, 1996) However, all sequenced chimeric clones, including cPOX04-12, carry the A34R gene from MVA, which lacks the K151 mutation. These findings suggest that additional genetic determinants beyond A34R may contribute to EEV production and comet formation in chimeric viruses, and their identification warrants further investigation. These findings underscore the potential of yeast-based TAR cloning as a powerful platform for generating diverse chimeric poxviruses. By enabling the shuffling of genomes from multiple parental strains, this approach provides access to a rich source of genetic and phenotypic diversity. Future studies leveraging high-throughput sequencing and phenotypic screening of larger chimeric virus libraries could facilitate genetic mapping of key traits such as syncytia formation, EEV production, oncolytic potency and immunogenicity. Indeed, identifying new genes responsible for EEV and syncytia formation could improve new oncolytic virus potential. And inversely, implementing the A56R and A34R gene mutations in our chimeric viruses could lead to a significant improvement of their oncolytic properties.

Moreover, expanding the platform to include poxviruses from other genera could further enhance the diversity and utility of engineered viruses. Ultimately, this strategy holds promise for the rational design of optimized oncolytic viruses capable of efficient tumor spread and immune evasion. One major advantage of our synthetic strategy for generating chimeric oncolytic poxviruses is the complete avoidance of *E. coli* in the cloning process. Cloning large and structurally complex viral genomes in *E. coli* using bacterial artificial chromosomes (BACs) presents well-documented technical challenges. For example, several groups have reported difficulties in cloning and manipulating coronavirus genomes due to their size and complexity, while high G+C content has been implicated in the cloning inefficiency of herpesviruses (Hall *et al*, 2012; Cao *et al*, 2023; Xiao *et al*, 2023; Knickmann *et al*, 2024). Moreover, bacterial systems are inherently limited in their capacity to support simultaneous manipulation of multiple genomic loci, and extensive modifications often prolong the timeline required to generate mutant viruses (Vashee *et al*, 2017; Oldfield *et al*, 2017). Instability of full-length viral cDNA clones in *E. coli* has been frequently observed for flaviviruses, herpesviruses, and coronaviruses, often due to toxicity induced by viral sequences during bacterial propagation (Polo *et al*, 1997; Kanda *et al*, 2011; Yakushko *et al*, 2011; Almazán *et al*, 2000). In some cases, cryptic bacterial promoters within viral genomes drive the expression of toxic proteins, as seen with Zika virus, where high-copy-number plasmids exacerbate this effect (Li *et al*, 2011). Our yeast-based synthetic approach circumvents these limitations, enabling efficient and stable assembly of complex viral genomes without bacterial propagation. This strategy represents a robust and scalable alternative for the generation of chimeric oncolytic poxviruses, facilitating rapid and precise genome engineering. Historically, Domi and Moss demonstrated the feasibility of cloning the VACV genome into a BAC, allowing stable maintenance and precise genetic manipulation of the full-length viral genome in *E. coli* (Domi & Moss, 2002). Their method relied on the accumulation of head-to-tail concatemers of VACV DNA, which were circularized and introduced into bacteria. This BAC-based system enabled efficient genome editing via recombineering and recovery of infectious virus through transfection into mammalian cells co-infected with a helper fowlpox virus (FPV). While well-suited for functional studies and complex genome engineering, this strategy requires careful control of concatemer formation and circularization, and the retention of bacterial sequences in the rescued virus may necessitate additional removal steps for translational applications. To overcome these limitations, *Chiuppesi et al.* developed a synthetic MVA platform that assembles the MVA genome from three synthetic BAC-cloned fragments and reconstitutes the virus via homologous recombination in permissive cells (Chiuppesi *et al*, 2020). This method enables rapid generation of recombinant viruses without plaque purification, and the spontaneous loss of bacterial sequences during virus rescue enhances its translational potential. Nevertheless, the system still relies on *E. coli* for BAC propagation, which may introduce bacterial elements or recombination artifacts. More recently, Gao et al. reported the complete synthetic assembly of the 180-kb MVA genome using 37 chemically synthesized DNA fragments and TAR technology (Gao *et al*, 2025). The resulting system, composed of five plasmids, enables rapid and modular construction of recombinant MVA vectors, significantly accelerating vaccine development. Despite its advantages, the method remains technically demanding and resource-intensive, and its reliance on *E. coli* for BAC propagation introduces potential concerns regarding plasmid stability and recombination fidelity. In conclusion, the generation of chimeric oncolytic viruses through synthetic recombination represents a promising avenue for cancer therapy. The phenotypic diversity observed among the clones—ranging from syncytia formation to differential EEV release and variable oncolytic potency—underscores the potential to tailor viral properties to specific tumor contexts. Notably, clones such as cPOX04-12, which exhibit enhanced activity in colorectal cancer cells and a pronounced comet phenotype, may be particularly advantageous for tumors with dense architecture, where long-range viral dissemination is critical. The yeast-based assembly platform employed in this study enables rapid and flexible engineering of viral genomes, effectively overcoming the limitations associated with bacterial cloning systems. This approach facilitates high-throughput generation and screening of chimeric viruses with optimized tropism, immunogenicity, and safety profiles. Future investigations may focus on identifying the genetic determinants underlying syncytia formation and EEV release, which could be harnessed to enhance intratumoral spread or modulate host immune responses. From a broader therapeutic perspective, these synthetic chimeric viruses hold potential for integration into combination regimens with immune checkpoint inhibitors, chemotherapeutics, or targeted agents (Yuan *et al*, 2024). Their modular design also permits the incorporation of transgenes encoding immunostimulatory molecules, further expanding their utility as multifunctional oncolytic platforms. Collectively, the strategy described herein lays the foundation for a new generation of customizable oncolytic viruses with improved efficacy and adaptability across diverse cancer types.

## MATERIAL AND METHODS

### >Cells and viruses

*Saccharomyces cerevisiae* VL6-48 (ATCC MYA-3666™, genotype: MATa his3-Δ200 trp1-Δ1 ura3-52 ade2-101 lys2 ψ⁺ cir°) was cultured in YPD medium containing 1% yeast extract, 2% peptone and 2% glucose, supplemented with 40 mg·L⁻¹ adenine, at 30 °C with agitation. Transformed VL6-48 strains were selected and maintained in synthetic dropout medium lacking tryptophan (−TRP, Takara Bio, Japan). For solid media, 2% Bacto™ Agar (Dutscher, France) was added.

*Escherichia coli* EPI300 (LGC Biosearch Technologies, USA) was used for DNA cloning and grown in LB broth (Sigma-Aldrich, USA), with or without agar, supplemented when required with chloramphenicol at 34 µg·mL⁻¹ and kanamycin at 50 µg·mL⁻¹ (Sigma-Aldrich, USA).

Human tumor cell lines A549 (ATCC CCL-185), HeLa (ATCC CCL-2), MIA-PaCa-2 (ATCC CRL-1420), HCT116 (ATCC CCL-247), and the African green monkey kidney cell line Vero (ATCC CCL-81) were maintained at 37 °C in a humidified atmosphere of 5% CO₂ in Dulbecco’s Modified Eagle Medium (DMEM, Thermo Fisher Scientific, USA) or McCoy’s 5A medium (LGC Standards, UK), supplemented with 10% fetal bovine serum (FBS, Thermo Fisher Scientific, USA) and 100 µg·mL⁻¹ gentamycin (Sigma-Aldrich, USA).

Wild-type Vaccinia virus strain Copenhagen (VACV) was obtained from Institut Mérieux (France), while wild-type Cowpox virus strain Brighton (CPXV; ATCC VR-302) and wild-type Rabbitpox virus strain Utrecht (RPXV; ATCC VR-1591) were obtained from ATCC (USA). VACV, CPXV and RPXV stocks were propagated in HeLa cells using standard procedures. Modified Vaccinia virus Ankara (MVA) (Spehner *et al*, 2000) was amplified in primary chicken embryo fibroblasts (CEF) as previously described (Erbs *et al*, 2008b).

### Molecular biology

The pYCC plasmid was constructed using the pCC1-FOS copy-control plasmid (LGC Biosearch Technologies, USA) as template. The yeast replication region CEN6-ARS4 and the autotrophic selection gene TRP1 were amplified as a single DNA fragment from the pRS314 plasmid (Sikorski & Hieter, 1989) by PCR using Primestar DNA polymerase (Takara Bio, Japan). The PCR product was inserted into pCC-FOS1 by InFusion® cloning (Takara Bio, Japan) at the AfeI restriction site (Hou *et al*, 2016). pCC-FOS1 was digested with AfeI (New England Biolabs, USA) according to the manufacturer’s instructions. Linker sequences 5’SU-L3’ were designed using Geneious software (see Supplemental Table 1) and synthesized by GeneArt (Thermo Fisher Scientific, USA). The pYCC3 plasmid was generated by InFusion® cloning of EcoRI-HF–digested pYCC (New England Biolabs, USA) with linker 5’SU-L3’ following the manufacturer’s protocol, and the resulting products were chemically transformed into EPI300. Correct insertion was verified by Sanger sequencing (Eurofins Genomics, Germany).

Viral DNA was extracted from 2 mL of purified virus using the following procedure: incubation for 2 h at room temperature with 500 units benzonase (EMD Millipore, USA) and 0.5 mM MgCl₂ (Sigma-Aldrich, USA); incubation for 2 h at 56 °C with 1 mg·mL⁻¹ proteinase K (Interchim, France); incubation for 2 h at 37 °C with 0.25 mg·mL⁻¹ RNase A (Qiagen, Germany). The extract was then sequentially treated with one volume of phenol (Interchim, France), centrifuged at 8000 rpm for 10 min, and the aqueous phase recovered; one volume of phenol–dichloromethane (1:1, Interchim, France) with centrifugation and recovery of the aqueous phase; and one volume of dichloromethane, followed by centrifugation and collection of the aqueous phase. DNA was precipitated for 2 h at −20 °C with two volumes of absolute ethanol (VWR International, USA) and 0.15 M sodium acetate (Sigma-Aldrich, USA), centrifuged at 8000 rpm for 30 min, and the pellet washed twice with ethanol/TE buffer pH 7.5 (7:3). DNA was resuspended in 200 µL of DNase-free water. For PCR screening of yeast clones, cultures were grown overnight at 30 °C in 50 µL SD-TRP selective medium with agitation, diluted with 100 µL water, and 5 µL of each suspension used as PCR template. Taq polymerase (Qiagen, Germany) was used with an initial denaturation step of 15 min at 98 °C (Bonnet *et al*, 2013). Primers are listed in Supplemental Table 2.

For plasmid DNA extraction, transformed EPI300 cultures were grown overnight in 150 mL selective medium supplemented with CopyControl Induction Solution (LGC Biosearch Technologies, USA). Transformed yeast cultures were grown for 48 h in 150 mL selective medium. DNA extractions were performed using Plasmid DNA Midiprep kits (Qiagen, Germany) and concentrates purified with NucleoSnap Finisher Midi (Macherey-Nagel, Germany) following manufacturer instructions. Prior to extraction, yeast cells were treated for 1 h with 800 µL zymolyase 20T and 80 µL of 14.3 M 2-mercaptoethanol (Sigma-Aldrich, USA).

EPI300 chemically competent cells were prepared by growing cultures at 37 °C in SOB medium to OD₆₀₀ ≈ 0.6, washing, and freezing at −80 °C in CCMB solution (10 mM potassium acetate, 80 mM CaCl₂, 20 mM MnCl₂, 10 mM MgCl₂, 10% glycerol in sterile water). TransfoMax EPI300 (LGC Biosearch Technologies, USA) was used for transformation of yeast-derived DNA. Electroporation was performed with 50 µL competent cells and 2 µL yeast DNA extract in 1 mm cuvettes using a Bio-Rad Gene Pulser II (USA) at 2.5 kV, 200 Ω and 25 µF. Restriction profiling of pYCC3-VACV plasmids was performed by digesting 500 ng of *E. coli* extract with SfiI (New England Biolabs, USA), followed by migration on a 0.8% agarose gel at 120 V for 1 h.

Restriction enzymes used in this study included EcoRI, AfeI, PacI, SphI and SfiI (New England Biolabs, USA) as well as SgrDI, MreI and FspAI (Thermo Fisher Scientific, USA).

### Yeast Spheroplast transformation

To prepare yeast spheroplasts competent for large DNA fragments, a fresh VL6-48 colony was inoculated into 20 mL YPDA and grown overnight, then used to inoculate 50 mL YPDA at an initial OD₆₆₀ of 0.4 and cultured until OD₆₆₀ reached 2–4. Cells were harvested by centrifugation at 1000 g for 5 min, washed once with 30 mL sterile water and once with 20 mL 1 M D-sorbitol, then resuspended and incubated for 25 min in 20 mL SPE solution (1 M D-sorbitol, 10 mM Na₂HPO₄, 10 mM EDTA) containing 0.1% zymolyase solution (1% zymolyase 20T, 50 mM Tris-HCl, 25% glycerol; Seikagaku, Japan) and 0.2% 14.3 M β-mercaptoethanol (Sigma-Aldrich, USA). After centrifugation at 500 g for 10 min, the pellet was washed twice with 30 mL of 1 M D-sorbitol and resuspended in 1 mL STC solution (1 M D-sorbitol, 10 mM Tris-HCl, 10 mM CaCl₂). For each transformation, 100 µL of spheroplast suspension was used.

For construction of pYCC3-VACV, DNA was mixed as follows: 0.5 µg AfeI-digested pYCC3, 1 µg VACV DNA digested with SgrDI, 1 µg digested with MreI, and 1 µg digested with FspAI (restriction enzymes from New England Biolabs, USA, and Thermo Fisher Scientific, USA). For viral shuffling, 0.5 µg PacI-digested pYCC3-VACV was mixed with 3 µg CPXV DNA digested with SphI and 3 µg RPXV DNA digested with PacI. Spheroplasts and DNA were incubated together for 10 min at room temperature, followed by incubation for 10 min with 800 µL PEG8000 solution (20% PEG8000, 10 mM CaCl₂, 10 mM Tris-HCl). After centrifugation at 500 g for 5 min, pellets were resuspended in SOS solution (1 M D-sorbitol, 0.65 mM CaCl₂, 0.25% yeast extract, 0.5% peptone) and incubated for 40 min at 30 °C without agitation. The suspension was mixed with 7 mL of melted selective medium cooled to 50 °C and poured onto selective plates, which were incubated for 5 days at 30 °C. Small patches of primary transformants were then transferred to SD-TRP agar plates and incubated for 2 days at 30 °C.

### Viral rescue

To restore the infectivity of viral DNA, 1 × 10⁵ A549 or HeLa cells were infected in 24-well plates at a multiplicity of infection (MOI) of 1 using MVA diluted in 200 µL of PBS 1× (Sigma-Aldrich, USA) supplemented with 1% FBS, 0.6% magnesium acetate and 0.75% calcium chloride, and incubated for 1 h at 37 °C in 5% CO₂ (Erbs *et al*, 2008a). Transfection of 500 ng yeast-extracted DNA was performed using the Lipofectamine 3000 kit (Thermo Fisher Scientific, USA) according to the manufacturer’s instructions. Plates were incubated for 7 days at 37 °C in 5% CO₂, and 500 µL of supplemented DMEM was added 24 h after transfection. After 7 days, the 24-well plates were frozen at −20 °C for 24 h, and cells together with their supernatant were harvested and sonicated for 30 s at 40% amplitude. A volume of 50 µL of each lysate was then used to infect 2 × 10⁵ A549 or HeLa cells in 6-well plates, followed by incubation for 3 days at 37 °C in 5% CO₂.

### Viral DNA extraction from infected cells

For viral DNA sequencing, 2 × 10⁵ A549 cells seeded in 6-well plates were infected with VACV or with an isolated viral clone in 250 µL PBS 1× supplemented with 1% FBS at 37 °C in 5% CO₂ for 30 min. After infection, cells were overlaid with supplemented DMEM and incubated for 3 days at 37 °C in 5% CO₂. Plates were then frozen at −20 °C, thawed slowly, and the resulting lysates were sonicated for 30 s at 40% amplitude. A volume of 100 µL of each infectious lysate was mixed with 5 mL supplemented PBS and used to infect an 80% confluent T175 flask of A549 cells for 30 min at 37 °C. Infected flasks were incubated for 3 days at 37 °C in 5% CO₂ and subsequently frozen at −80 °C. Thawed cell suspensions were centrifuged for 3 min at 1000 g, the supernatant discarded, and the pellet resuspended in 9 mL PBS 1×. Viral DNA was extracted from 1 mL of this suspension using the Monarch HMW DNA Extraction Kit (New England Biolabs, USA) according to the manufacturer’s instructions.

### Cytotoxicity assay

The lytic capacity of viruses was assessed using the Trypan blue exclusion method. HeLa, A549, HCT116 and MIA PaCa-2 cells were infected in suspension with serial virus dilutions corresponding to multiplicities of infection (MOI) ranging from 3 × 10⁻⁵ to 1, after which 1 × 10⁴ cells per well were seeded into 96-well plates in 150 µL of supplemented DMEM. Infected cells were incubated for 4 days at 37 °C. Cell viability was then quantified using the CellTiter-Blue assay (Promega, USA), and fluorescence readings were obtained using a Spark® multimode microplate reader (Tecan, Switzerland). All conditions were analyzed in triplicate, and non-infected cells were used as negative controls to establish the 100 % survival reference for each assay.

### Syncytia formation assay

HeLa cells were infected with the indicated viruses at a multiplicity of infection (MOI) of 10⁻² or 10⁻³, and 3 × 10⁵ cells per well were seeded into 6-well culture dishes. Infected cells were incubated at 37 °C in a humidified 5% CO₂ atmosphere and examined by optical microscopy 72 h post-infection to assess syncytia formation.

### Comet assays

A549 cells were seeded into 60 mm tissue culture dishes and grown to confluence overnight. The following day, monolayers were rinsed with PBS and inoculated with approximately 40 plaque-forming units (PFU) of the indicated viruses in 500 µL PBS supplemented with 2% FCS and 1% cations, followed by incubation for 1 h at 37 °C. After adsorption, monolayers were rinsed and overlaid with 5 mL growth medium. Two days post-infection, the cell monolayers were fixed and stained with crystal violet (stock solution diluted 1:40 in ethanol) to visualize viral plaques.

### *In vitro* virus yield

To evaluate viral replication in human tumor cells, A549 cells were infected in 6-well plates at an MOI of 10⁻³ in triplicate. Three days post-infection, both supernatants and cell fractions were collected, subjected to one freeze–thaw cycle, sonicated, and the viral progeny was quantified on Vero cells by plaque assay. To quantify infectious extracellular enveloped virions (EEV) and intracellular mature virions (IMV) at early time points, confluent A549 cells (1 × 10⁶ cells) were infected in 6-well plates at an MOI of 10⁻¹ for 1 h in triplicate, washed with PBS, and overlaid with fresh medium before incubation for 24 h at 37 °C. Supernatants were collected to quantify EEV, whereas cell-associated virions (IMV) were recovered by resuspending the cell pellets in 1 mL PBS, followed by freeze–thawing and sonication. Both EEV and IMV fractions were titrated on Vero cells by plaque assay.

### DNA sequencing

Viral DNA samples extracted using NEB Monarch HMW DNA Extraction Kit were internally sequenced using MinION™ (Oxford Nanopore Technologies®) for long reads sequencing data analysis. The raw data files from R10.4.1 MinION flow cells were transferred from the lab to a high computation workstation. A data processing workflow, developed in-house to ensure reproducibility and automation of the analysis steps, has then been applied using a custom Snakemake pipeline. The steps are described hereafter. The raw electric signals (pod5 files) are converted into nucleic acid bases (fastq files) using guppy_basecaller software (v6.5.7) with dna_r10.4.1_e8.2_260bps_sup.cfg model file. The demultiplexing step was performed with guppy_barcoder (v6.5.7) to affect specific long reads to the respective sample. A quality control with pycoQC (v2.5.2) generates metrics from the sequencing run (number of reads, median read length, quality PHRED scores, barcode proportions, …). For each sample, reads are trimmed from the adapters with porechop (v0.2.4) and are filtered using NanoFilt (v2.8.0) with a minimum average PHRED quality score of Q12. Canu (v2.2) was then used for the de novo assembly from the filtered reads in combination with Medaka (v1.8.0) for the final polishing step. The contigs from each sample were imported in the Geneious software (v.8.1.9) for visualization and sequence alignments against the theoretical reference sequence. The filtered reads have then been mapped with minimap2 on the longest contig of each sample to check the variations (single nucleotide variants, insertions, deletions) by visual inspection with IGV (v2.11.9).

### Statistical analysis

Data were analyzed using one-way analysis of variance (ANOVA) followed by post-hoc comparisons with Tukey’s adjustment. Differences were considered statistically significant at p < 0.05, with significance levels indicated as follows: p < 0.05 (*) and p < 0.01 (**).

## Author contributions

A. Agaoua and P. Erbs conceived the project and wrote the manuscript. A. Agaoua, C. Rey, J. Hortelano, A-I. Moro, B. Grellier and P. Erbs designed, performed and analyzed the experiments.

## Acknowledgements

We thank Anita Spinler, Karine Reymann, and Véronique Koerper (Transgene) for virus production and viral genome extraction. We thank Delphine Suhner (Transgene) for sampling viral DNA for DNA sequencing. We thank Berangère Bastien (Transgene) for the statistical analysis.

